# Spatial Variation of Soil Moisture and its Effect on Lint Yield in a Deficit Irrigated Cotton Field

**DOI:** 10.1101/109025

**Authors:** Yujin Wen

## Abstract

As the population increases in Southwest Texas in recent years, the urban water demand is drastically increasing. Regulated deficit irrigation (RDI) is expected to be one of the potential water management practice for saving water while maintaining crop yield. A field experiment was conducted at the AgriLIFE Research center in Uvalde in summer 2008 to examine the water saving potential. Seven irrigation schemes and four varieties were assigned to the experimental field to test their effects on lint yield. As the spatial correlation of the soil moisture/ soil water content were suspected, a spatial analysis on lint yield and soil water content was conducted. The analysis results showed that: 1) The soil water contents showed spatial correlations, while the lint yield did not. 2) The relationship between lint yield and soil water content can be described by linear model better than spatial autocorrelation model. 3) The variogram fit of the soil water content showed a completed curve; the contour map generated using ordinary kriging illustrated the irrigation schemes effect well, and gradient effect was suspected. The variogram of the lint yield could not be fitted by a completed curve, which indicated that the sample field was not large enough to determine the variance function. Further study is needed to determine the slope effect on soil water content, and to improve the contour map precision of the lint yield.

## INTRODUCTION

As the population increases in Southwest Texas in recent years, the urban water demand is drastically increasing. Since the water resources in this area are limited, making a good plan for the available water supply is crucial. One possible way to assist in solving this problem is to reduce the agricultural water use; however, the economic crop yield, or the growers' profit, should at least be maintained. Regulated deficit irrigation (RDI) is one important measure for saving water and maintaining crop yield. A field experiment was conducted at the AgriLIFE Research center in Uvalde in summer 2008 to examine the water saving potential (Wen et al., 2009, 2013). Since the spatial correlation of the soil moisture/ water content, which may cause lint yield variation (Ge et al., 2008; Johnson et al., 2002), is suspected, it is essential to test the soil spatial correlation before the other data analysis procedure. The spatial variation of lint yield is also of interests.

Many studies discussed the spatial analyses methods for soil moisture and agronomic yields, such as Bi et al. (2008) and de Lannoy et al. (2006). The objectives of this study are: 1) find out whether the soil water content and the lint yield data are spatially correlated; 2) if correlations are found, whether the lint yield can be predicted by the soil water content via a spatial model; 3) test the spatial variation of lint yield and soil water content and build contour maps to check whether the spatial correlation was caused only by irrigation, or also by some other factors.

## MATERIALS and METHODS

### The Cotton RDI Experiment in 2008

The research was conducted in the AgriLIFE Research and Extension Center at Uvalde in summer 2008 (Wen et al., 2009, 2013). A split-split-plot design experiment was assigned in a 90^o^ wedge (approximate 40 ac) (Wen, 2015). The wedge was divided into four spans (160 ft width each) and two "buffer zones" (filler spans). Each span had 48 rows, which were divided into four 12-row zones. Four widely planted commercial varieties were randomly selected, which were DP555, DP164, FM9063, and 989B2R. These four varieties were randomly assigned to the four zones in each span. The cotton was planted on April 15, 2008, and harvested on September 25 (162 day after planting /162 DAP)).

Irrigation was applied by a center pivot with a low energy precision application (LEPA) system with 95% efficiency. Seven irrigation regimes were selected, including the fully irrigation (100X, as the control), four fixed deficit irrigation: 80X, 70X, 60X and 50X, and two dynamic irrigation: 70D and 50D. The irrigation replacement is described in percentage of the net evapotranspiration (ΔET), which equals to the difference between evapotranspiration (ET) and rainfall (P) in a certain period:

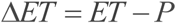

For instance, the number 50 in 50X stands for 50% replacement, that is, for each 1 mm water loss in the net evapotranspiration, we provide 0.5 mm water back to the field through irrigation. In practice, we recorded the daily ET and P to calculate the daily net water loss (ΔET), and then accumulated the net water loss day by day until it reaches to a certain maximal limit (we used 3.8 mm (1.5 in)), at which we applied irrigation. In the fixed scheme (marked as X), the replacement rate (as percentage of ΔET) is kept constant, e.g. 50X means in each irrigation application, we compensated the field with 50% of the water loss. In the dynamic scheme (marked as D), the irrigation is applied in different replacement ratios at each growing stages, for instance, in this study 50D was scheduled as 50% at vegetative stage, 100% from first bloom to 50%-open boll, and 10% thereafter till harvest. In total, the water use should be maintained at a certain range between 45% and 55% (depends on how much rain we receive at each growing stage) for 50D irrigation scheme.

### The Data Collection and Analysis

The soil moisture at 20cm, 40cm, 60cm, 80cm, 100cm, 120cm and 140cm were measured seven times (Jun.19, Jun. 26, Jun. 30, Jul. 10, Jul. 18, Jul. 28, and Aug. 5) using two neutron probes (CPN-530 DR Hydroprobe Probe Moisture Depth Gauge, Campbell Pacific Nuclear Corp. Int. Inc., Martinez, CA) during the cotton growing season. After planting, neutron probe access tubes were installed at the center of each experimental unit. Volumetric water content of each layer was calculated using the calibrated linear equations (one equation per layer), and the soil water content on each experimental unit in each measurement was derived. On September 25, 12 m^2^ areas were randomly selected in each experimental unit, and all seed cotton were harvested in these sample areas by cotton picker. Then small samples were selected from each harvested sample, and ginned in the Cotton Improvement Lab (Texas A&M Univ., College Station, TX). According to the weight ratio of lint to seed cotton of the small samples, the lint yield in each experimental unit was estimated. The coordinates of the centers of each experimental unit were calculated based on the distances to the center pivot engine and the angles to the North direction (Fig. 1).

**Figure 1:**
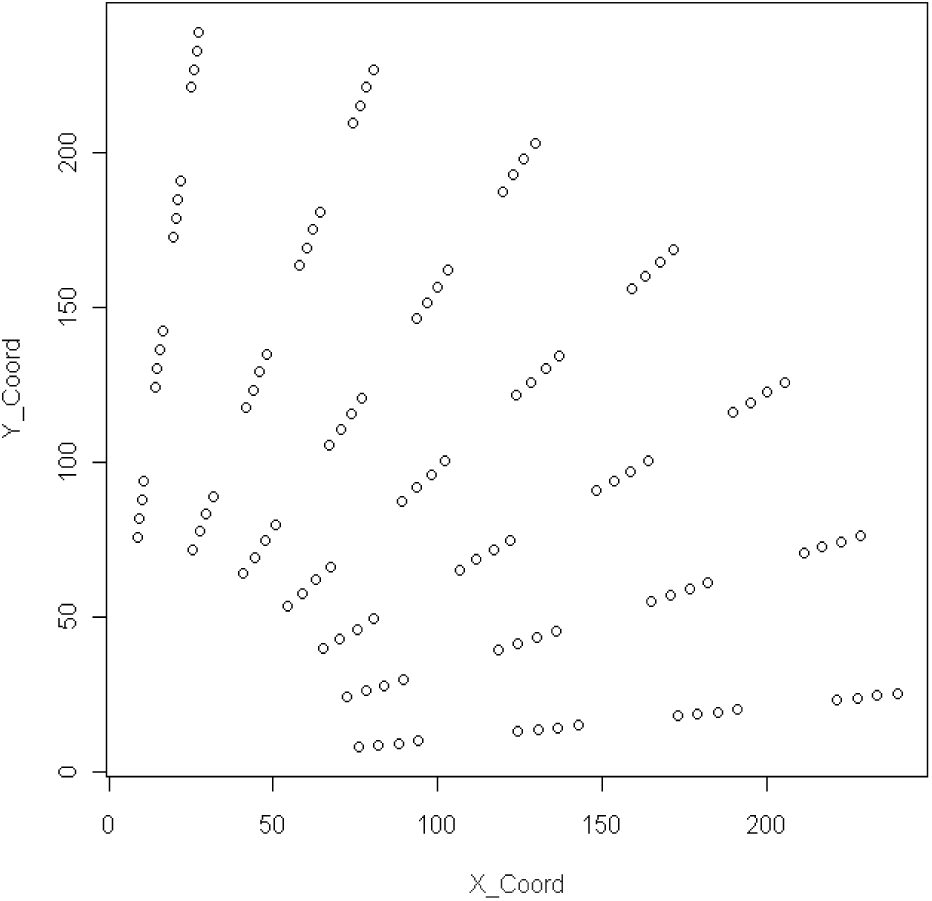
The positions of the sample points in each experimental unit

The data were first tested by the inverse-distance weighted Moran's I to check the possible spatial correlation. Possible models of lint yield vs. soil water content were fitted and compared to select the best one that can describe the relationship between them. The variograms of the lint yield and soil water content were calculated based on the sample data, and the contour maps were derived using ordinary kriging. All the spatial analysis were done using spdep and geoR packages installed in R (2.8.1). The default 0.05 significance level was used in this study.

## RESULTS and DISCUSSIONS

### Determine the Spatial Correlations

The Moran's I test statistics were calculated for each variable, showed in Table 1. W1 to W7 are the soil water content at seven different times. The results indicated that the soil water content at the 4^th^ measurement did not have spatial correlation. Neither did the lint yield. The other soil water content variables had spatial correlations. According to the irrigation-rainfall records, we had relatively lower ET and a bit rainfall right before the 4^th^ measurement, which may cause lower water consumption by plants, thus the soil water content differences among irrigation schemes did not show strong spatial correlation. In general, soil water content had spatial correlation, while lint yield did not.

**Table 1:** **The inverse-distance based Moran's I test**

| Variable | Moran's I | Standard Dev. | P-value | Significance |
| --- | --- | --- | --- | --- |
| W1 | 0.6168 | 8.1342 | 0.0000 |  |
| W2 | 0.3239 | 4.3343 | 0.0000 |  |
| W3 | 0.1870 | 2.5459 | 0.0059 |  |
| W4 | 0.0808 | 1.1629 | 0.1224 | NS* |
| W5 | 0.3586 | 4.7615 | 0.0060 |  |
| W6 | 0.2805 | 3.7648 | 0.0000 |  |
| W7 | 0.3650 | 4.8767 | 0.0000 |  |
| LintYield | 0.0523 | 0.8062 | 0.2101 | NS |

\* NS: not significant. In this case it means no spatial correlation.

### Model Fits and Comparison

The linear relationship between lint yield and soil water content were fit by several different models and the fit results were compared (Tab. 2). For both linear regression and SAR models, different combinations of soil water content measurements were tried and all non-significant terms were removed, except the intercept. Both linear regression and SAR models agreed that the 3^rd^ soil water content measurement was the best predictor for lint yield. The difference between LM1 and LM2 was LM2 did not include intercept, which might be a better model comparing to LM1. The AIC of SLM1 indicated that SLM1 is not significantly better than LM1 (1667.4 > 1665.5), and the non-significat lambda implied that the spatial autocorrelation is really weak; the LM2 did not show better fit than LM1, either (Tab. 3). Thus, LM1 seems to be the best model to predict lint yield in this case.

**Table 2:**
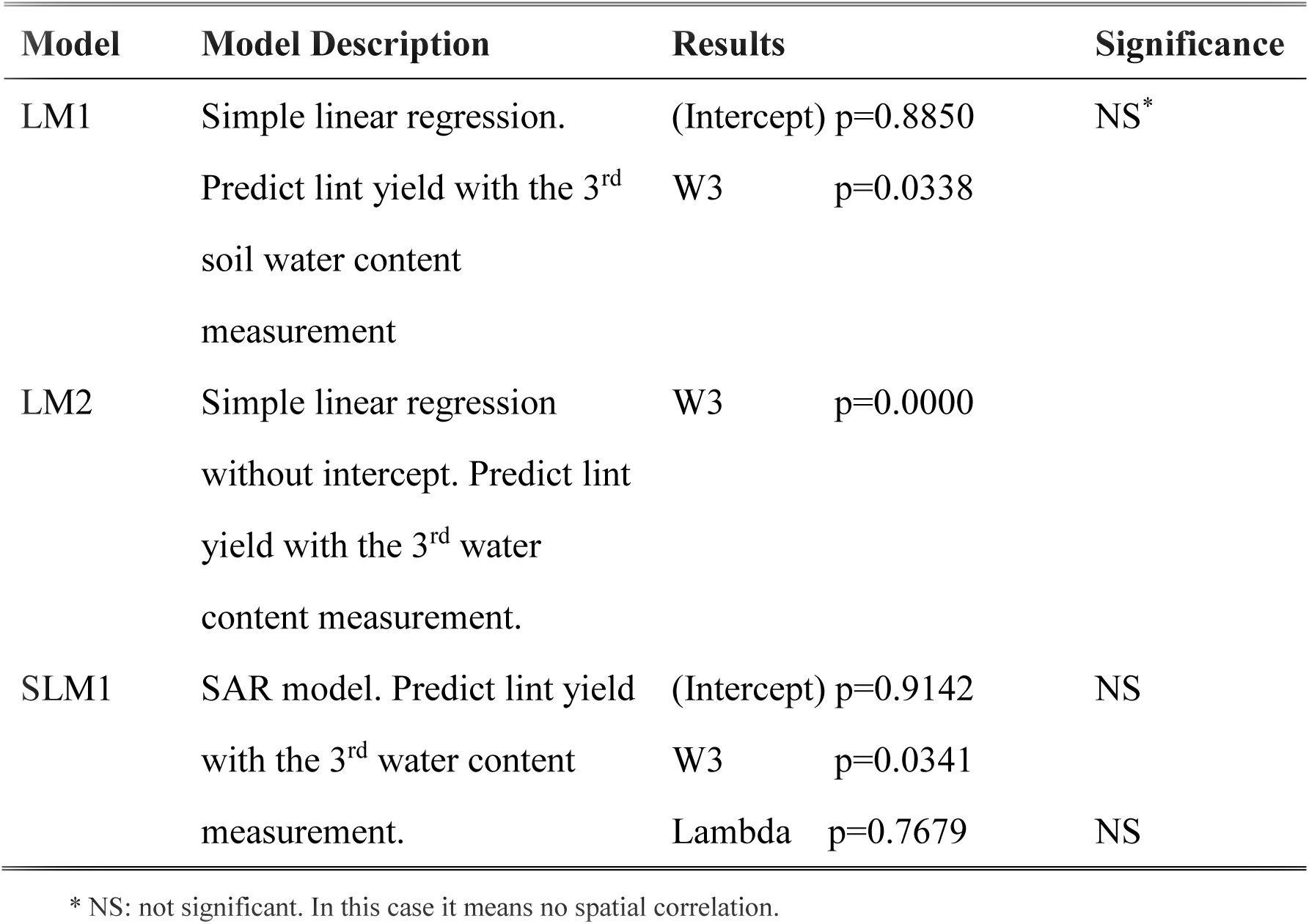
**Model-fit results of simple linear regression and SAR**

**Table 3:**
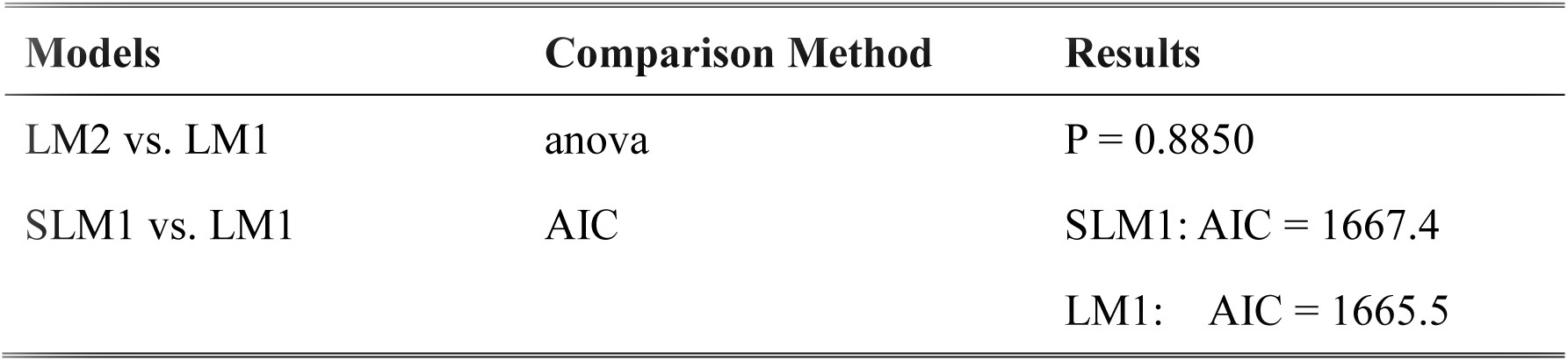
**The linear regression and SAR models comparison**

| Models | Comparison Method | Results |
| --- | --- | --- |
| LM2 vs. LM1 | anova | P = 0.8850 |
| SLM1 vs. LM1 | AIC | SLM1: AIC = 1667.4 |
|  |  | LM1: AIC = 1665.5 |

By examining the residuals of LM1 (Fig. 1), although the residuals demonstrated some curvature (left-hand side figure), it seems that the assumption of normal residuals is acceptable. Some potential outliers (e.g. #3, 4, and 87), as illustrated in Fig. 2, needed to be further examined though.

**Figure 2:**
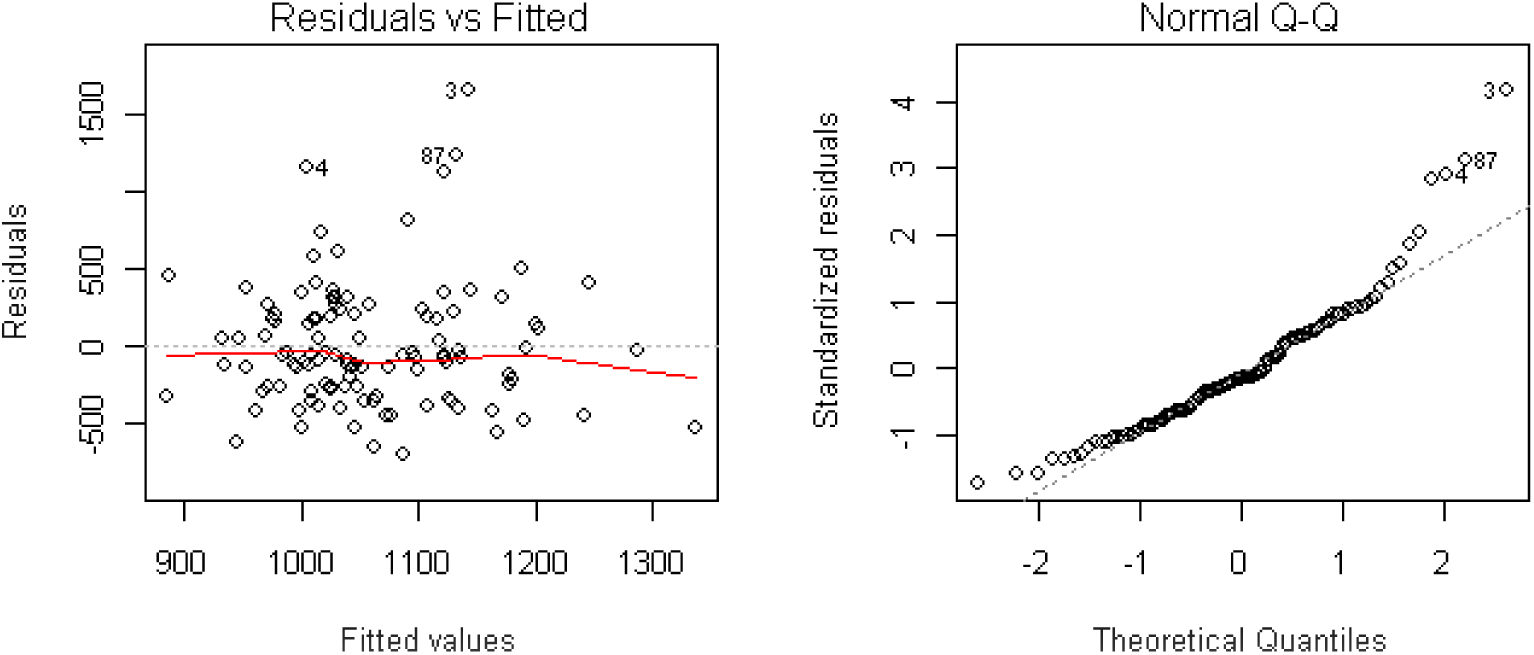
The diagnostic of LMI residuals. The left-hand side figure showed the residuals vs. fitted values. The right-hand side figure illustrated the Normal Q-Q plot.

In general, we concluded that the simple linear regression is the best fit for the lint yield and soil water content (3^rd^ measurement) relationship. Some other spatial model, e.g. Conditional Autoregression Model (CAR model), might be able to give a better prediction. Another concern is, the last soil water content measurement was not the best linear predictor of the lint yield. The reason why the best linear predictor is the 3^rd^ soil water content measurement is still unclear, thus further examine and more sampling on the soil moisture may be needed in the future study.

### Spatial Maps of the Lint Yield and Soil Water Content

The empirical variograms of the lint yield and the 3^rd^ soil water content measurement were calculated through the sample points and the fitted exponential models are shown in Fig. 3. The variogram fit of lint yield illustrated a linear trend, which indicated that the variance is very high, and the samples were not representative for the whole field. The variogram fit of W3, on the contrary, showed a reasonable variance and a range of approximate 40 m. A summary of the variogram fits was given in Table 4.

**Figure 3:**
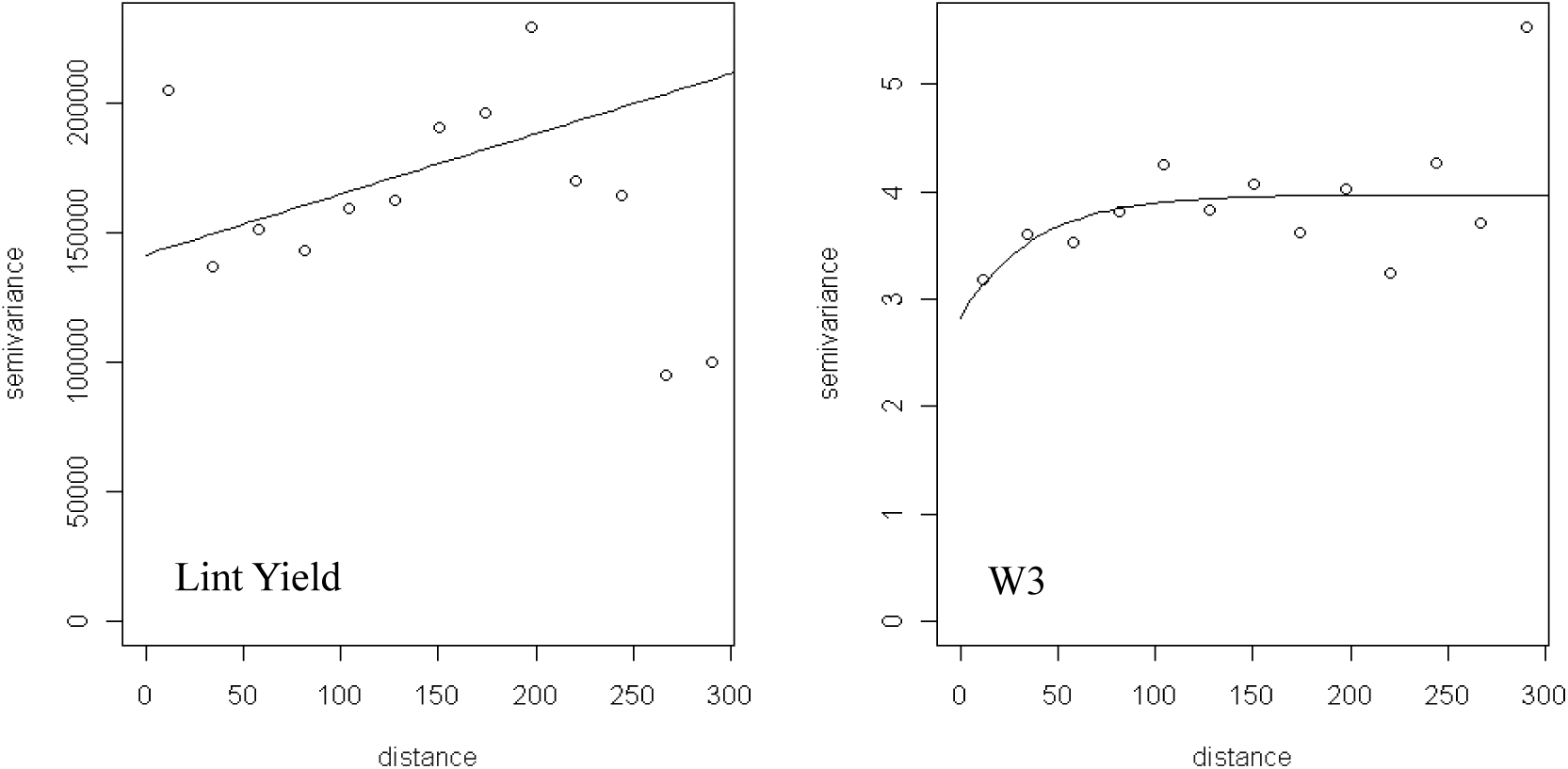
The variograms of lint yield and W3

**Table 4:** **Summary of the variogram fits of lint yield and W3.**

|  | Range [m] | Nugget | Sill | Nugget percent |
| --- | --- | --- | --- | --- |
| Lint Yield | 18242.39 | 141331.34 | 44259821.94 | 3.19% |
| W3 | 37.35 | 2.82 | 3.97 | 71.03% |

Based on the variogram fit functions, we plotted the contours of the lint yield and W3 (water content) use ordinary kriging. As the varigram fit function of lint yield did not show a completed curve, the kriging map could not show a very good spatial distribution of the lint yield (Fig. 4). The soil water content map demonstrated a good spatial distribution, confirming that the region that less water was applied has less soil water content left after a few days plant consumption. A gradient is shown as well, indicating that the soil water content, or soil moisture may be affected by slope. Further study on the slope or aspect effects need to be considered in the future study.

**Figure 4:**
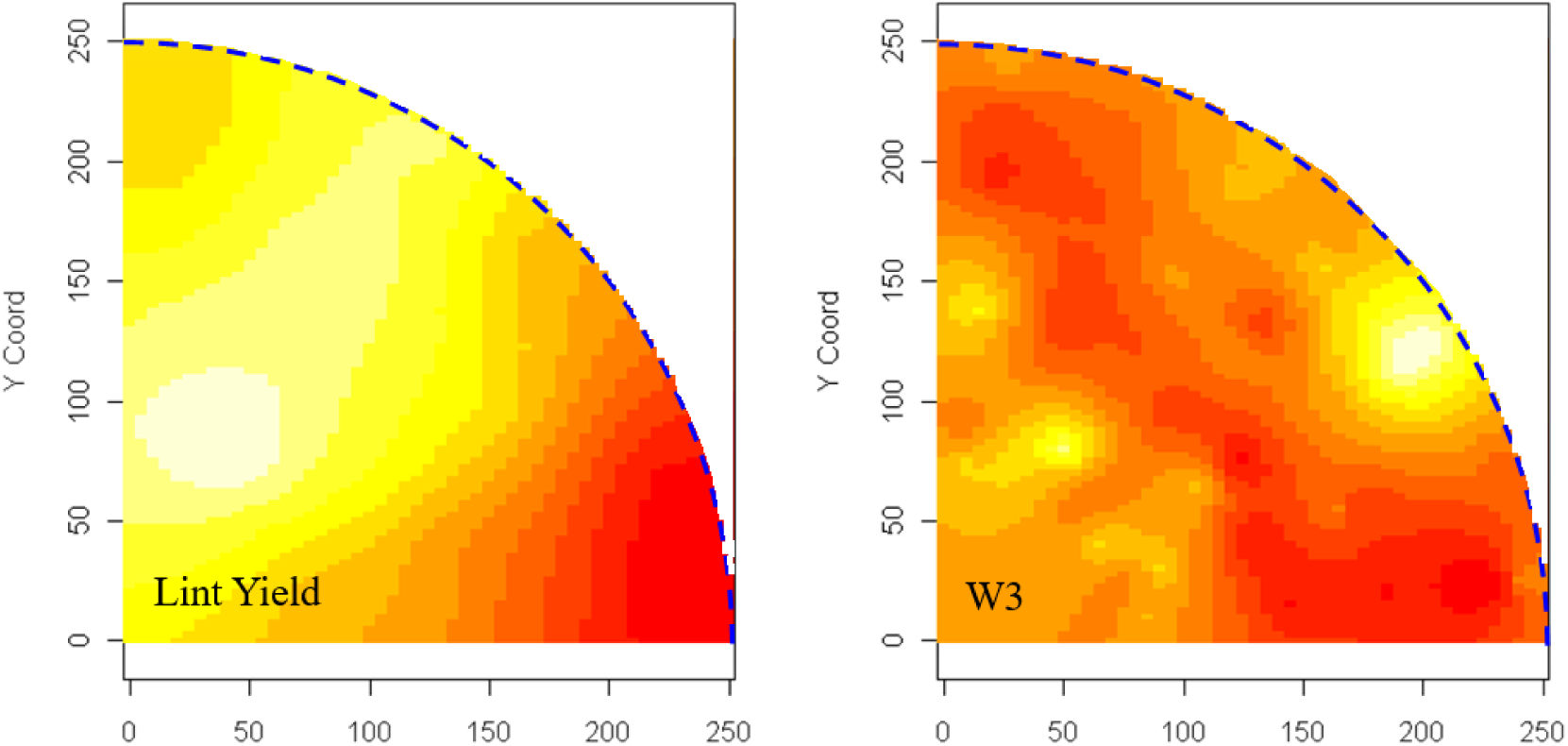
Contour maps of lint yield and W3 by ordinary kriging.

## CONCLUSIONS

Based on the results and discussion, the following conclusions can be drawn:

1. The soil water contents showed spatial correlations, while the lint yield did not.
2. The relationship between lint yield and soil water content can be described by linear model better than spatial autocorrelation model.
3. The variogram fit of the soil water content showed a completed curve; the contour map generated using ordinary kriging illustrated the irrigation schemes effect well, and gradient effect was suspected. The variogram of the lint yield could not be fitted by a completed curve, which indicated that the sample field was not large enough to determine the variance function. Further study is needed to determine the slope effect on soil water content, and to improve the contour map precision of the lint yield.

